# Experimental tests of functional molecular regeneration via a standard framework for coordinating synthetic cell building

**DOI:** 10.1101/2021.03.03.433818

**Authors:** Eric Wei, Drew Endy

## Abstract

The construction of synthetic cells from lifeless ensembles of molecules is expected to require integration of hundreds of genetically-encoded functions whose collective capacities enable self-reproduction in simple environments. To date the regenerative capacities of various life-essential functions tend to be evaluated on an ad hoc basis, with only a handful of functions tested at once and only successful results typically reported. Here, we develop a framework for systematically evaluating the capacity of a system to remake itself. Using the cell-free Protein synthesis Using Recombinant Elements (PURE) as a model system we apply our framework to evaluate the capacity of PURE, whose composition is completely known, to remake 36 life-essential functions. We find that only 23 of the components can be well tested and that only 19 of the 23 can be remade by the system itself; translation release factors remade by PURE are not fully functional. From both a qualitative and quantitative perspective PURE alone cannot remake PURE. We represent our findings via a standard visual form we call the Pureiodic Table that serves as a tool for tracking which life-essential functions can work together in remaking one another and what functions remain to be remade. We curate and represent all available data to create an expanded Pureiodic Table in support of collective coordination among ongoing but independent synthetic cell building efforts. The history of science and technology teaches us that how we organize ourselves will impact how we organize our cells, and vice versa.

## Introduction

Cells are the fundamental units of life, capable of growth, division, morphogenesis, and more. Yet, there is no natural cell for which we understand all of its life-essential functions [1]. Our lingering collective ignorance has practical costs that are difficult to quantify and overstate. For example, most of biotechnology research and development has remained an Edisonian process, grounded in trial and error, with state-of-the-art commercial platforms representing their prowess with automation or ‘atheoretic’ approach. Fully-distributed biomanufacturing, in which cells are optimized to operate reliably in many different local environments, remains practically impossible. As a second example, biosecurity strategies presume that we will always remain at risk from biology [2], a reasonable position given that each now-unknown life-essential function is a potential vulnerability to be exploited by the next natural emerging infectious disease, let alone by any mal-intentioned actor.

Understanding simple living cells in their entirety has been a dream of modern biology for almost 60 years [3]. The best ongoing examples are pioneering efforts to fully understand simple *mycoplasma* and *mesoplasma*. For example, sequencing followed by transposon mutagenesis, comparative and experimental genomics, and ultimately genome design and synthesis have been carefully combined to determine well-defined gene sets encoding all necessary and sufficient life-essential functions [4-9]. Practically, these approaches starkly reveal that dozens to hundreds of genes, each encoding life-essential functions, remain to be fully understood. Ongoing work to apply modeling and computation in guiding future experiments [10-11] suggests that many mysteries remain.

As a complementary approach scientists have long wondered about and explored the origins of life, both in the context of Earth’s history and beyond [12]. More recently the possibility of building simple cells from scratch has motivated the formation of world-leading scientific consortia; significant work to establish and benchmark the performance of individual life-essential functions is underway [13] (Fig. 1A-B). However, less well advanced is the consideration of challenges that loom inevitably for any bottom-up cell building effort. Most abstractly, just as natural genome minimization efforts highlight the mysteries of life-essential genes encoding unknown functions, bottom-up synthetic cell building efforts seem likely to encounter the puzzles of missing life-essential functions that are unknown (e.g., everything thought to be necessary and sufficient for life has been added but the hoped-for synthetic cell still does not grow and divide).

**Figure 1.**
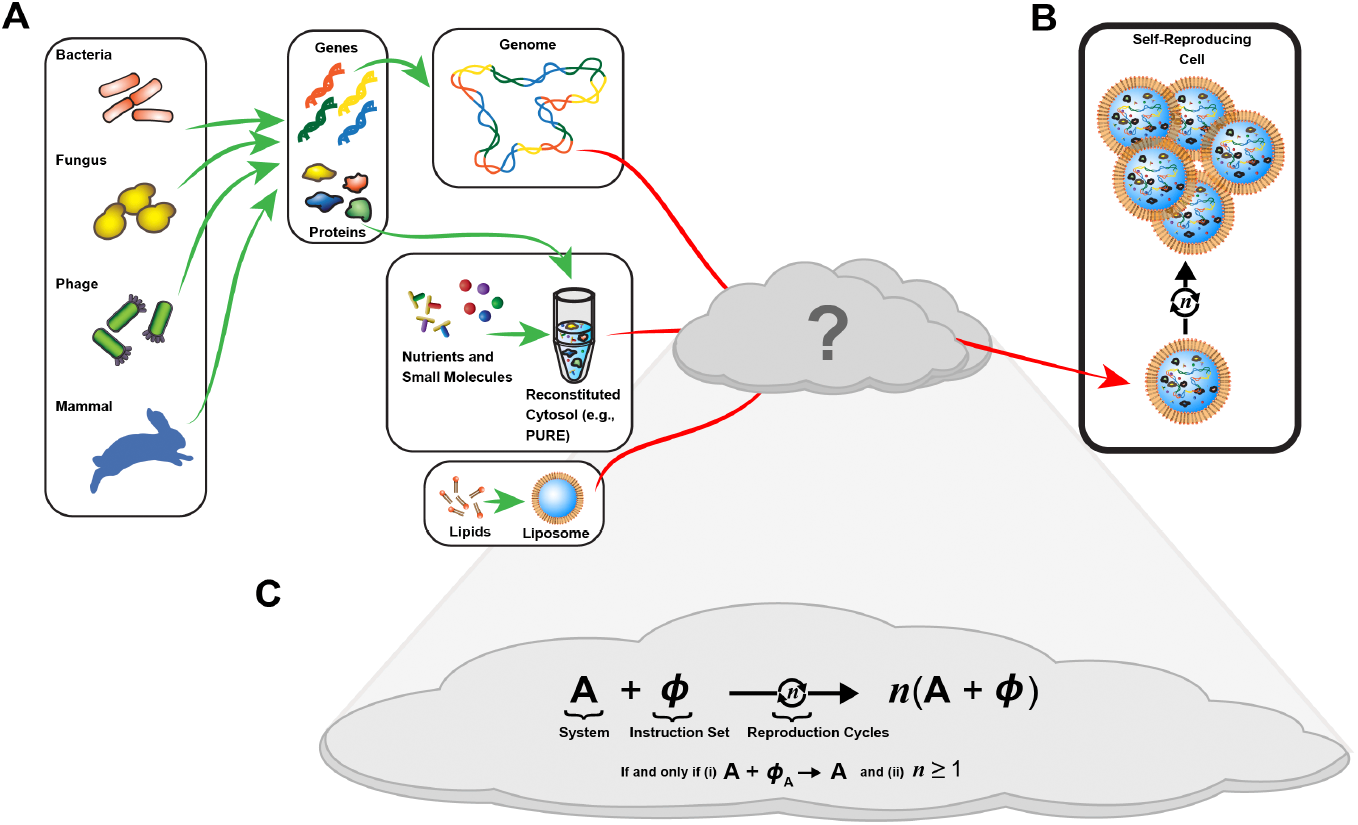
Can we build cells from lifeless ensembles of independently-sourced natural biomolecules? **(a)** We have the capacity to source, encapsulate, and encode the molecules thought to be essential for cellular functions. **(b)** Success would result in the capacity to produce autonomous reproducing simple cells comprised only of known components. **(c)** However, success requires that two abstract conditions be met. First, the ensemble itself must be capable of regenerating the functionality of all components comprising the ensemble (i.e., qualitative reproduction). Second, given energy and materials, the ensemble must be capable of remaking more of itself (i.e., quantitative growth).

From a first-principles perspective the engineering of physical systems capable of self-reproduction [14-15] must satisfy two criteria (Fig. 1C). First, qualitatively, the system enacting the instructions for reproducing the system must be capable of producing a functional copy of the system at both the level of individual components and as an integrated whole. Second, quantitatively, the system must possess sufficient generative capacity to make or organize at least as much material as needed to reproduce. For bottom-up synthetic cell building efforts the qualitative criteria means that the DNA encoding the system, when expressed in the environment defined within synthetic cell alone, must result in fully functional molecules for all so-encoded molecules. Next, satisfying the quantitative criteria means, for example, that the total number of peptide bonds used to instantiate proteins comprising the system can be catalyzed by the system within a single reproduction cycle.

The Protein synthesis Using Recombinant Elements (PURE) system enables transcription and translation of user-defined DNA [16]. PURE itself is well established as a research tool and benefits from reliable commercial and informal supply chains [17-18]. Many have imagined PURE or its analogs as a compelling biomolecular foundation upon which to build synthetic cells [19-20]. Practically, for PURE to form the basis of any free-living synthetic cell, PURE alone must satisfy the qualitative and quantitative criteria stated above or, more reasonably, when supplemented with additional functions. Stated differently, if we added the DNA encoding all of PURE to a mixture that began as PURE alone, would we get more working PURE made by PURE?

Simple calculations suggest that PURE itself cannot satisfy the quantitative criteria for self-reproduction. For example, on a quantitative volumetric basis, we estimate that PURE is only capable of remaking ∼1% of the peptide bonds needed to instantiate PURE itself; expression capacity will need to be increased ∼87-fold to enable sustained self-reproduction (Materials and Methods). However, such quantitative challenges can likely be addressed via exogenous supply of resources directly from the environment, via booting up of a regenerative and auto-sustaining metabolism, or by addition of enzymes and mechanisms that increase the efficiency of protein expression by reducing the number of truncated peptides produced and decreasing the rate of ribosomal stalling [17,21-25].

Whether PURE can satisfy the qualitative criteria for self-reproduction is less clear. PURE is well known to enable expression and synthesis of arbitrary genes, including many of the genes encoding PURE itself [25-26]. However, the molecules comprising PURE are made by and purified from intact cells, within natural contexts defined by the entire native complement of biochemical functions. Functional modification of one or more of the molecules comprising PURE via functions present in the source-cells may contribute to PURE activity yet not be present within PURE. As examples, Li et al. reported that *E. coli* ribosomes consisting of PURE-synthesized 30S r-proteins, native 16S rRNA, and a native 50S ribosomal subunit only had ∼13% activity compared to the native *E. coli* ribosomes [27]; Hibi et al. showed that PURE itself could simultaneously remake 21 tRNA, one for each corresponding amino acid and initiator tRNA, and when used by PURE recovered ∼40% of full PURE activity [28]; Lavickova et al. showed that PURE could regenerate the T7 RNA polymerase (RNAP) and eight tRNA synthetases within a diluting fluidic system initiated by functional PURE [29]; and Libicher et al. verified that PURE, through serial transfer, could make functional T7 RNAP, two energy recycling factors, and an ensemble of 12 tRNA synthetases and RF1 [30]. Separately, Libicher et al. and Doerr et al. used multi-plasmid systems to encode ∼30 of the translation factors of PURE and verified successful co-expression via mass spectrometry, but it is unclear whether these PURE-made enzymes were functional [25,31].

Here, we sought to develop and apply a method by which each molecule comprising a candidate auto-reproductive biomolecular system can be unambiguously evaluated in terms of the qualitative ability of the system to remake the molecule. We also wished to develop an approach by which additional molecules could be evaluated and accounted for. Finally, given the complexity of bottom-up synthetic cell building efforts and mindful of couplings between technical approach and cultural practice, what we sought was not just a method in which to evaluate molecules but a framework around which a fellowship could form. We know from the experience of other technologies that the essence of communal biotechnology will not only be about performing experiments in laboratories but to encourage close communication and collaboration [32].

## Results

We first sought to systematically determine and quantify which individual enzymatic components of PURE, when removed from PURE, result in diminished PURE activity as measured by expression of a reporter gene. Since the standard commercially-available starting material (PURExpress) combines all of the enzymes into a single tube (PURExpress Solution B), we produced our own custom mixtures consisting of single-enzyme depletions by purifying each enzyme comprising PURE and reconstituting single-enzyme depletion PURE variants.

We started by purifying all 20 tRNA synthetases (aaRSs) from *E. coli* (Fig. S2). We then tested whether the purified synthetases were functional by adding them, all together, to the PURE Solution B that lacked its own aaRS set and measuring expression of a green fluorescent protein (GFP) (Fig. S4A, Fig. S3, red curve). We chose GFP as a simple, easily accessible reporter for our study (Table S1). PURE Solution B lacking any aaRSs produced almost no GFP whereas GFP was well-expressed by PURE Solution B containing commercially supplied aaRSs (Fig. S3). We observed that the expression profile of PURE using our newly-purified aaRSs nearly matched those of the commercial mixture, suggesting that our so-purified aaRSs were all functional and could be used to construct each of the 20 aaRS single-depletion PURE mixtures.

To construct the remaining 16 single-enzyme depletion PURE variants we sourced six different custom PURE Solution B kits (NEB), each with a different subset of missing enzymes (Fig. S4). We then made all remaining single-enzyme depletion PURE variants by supplementing each depletion subset with the appropriate combinations of enzymes that we had independently purified from *E. coli* or sourced commercially (Methods).

We next measured GFP expression capacity for each single-component depletion (SCD) (Fig. 2, Fig S6-7). We use PURE_△ (component)_ to indicate specific SCDs. We expected some decrease in expression for all SCDs since each component tested is necessary for gene expression, in general, and our reporter gene (GFP) requires at least one of each of the 20 standard amino acids (Table S1). For example, PURE_△ (AlaRS)_(i.e., PURE lacking the alanine tRNA synthetase) produced ∼98% less in GFP expression relative to intact PURE (Fig. 2AB). However, we observed that each SCD produced a different level of GFP expression, from no change to almost complete loss of expression (Fig. 2C).

**Figure 2.**
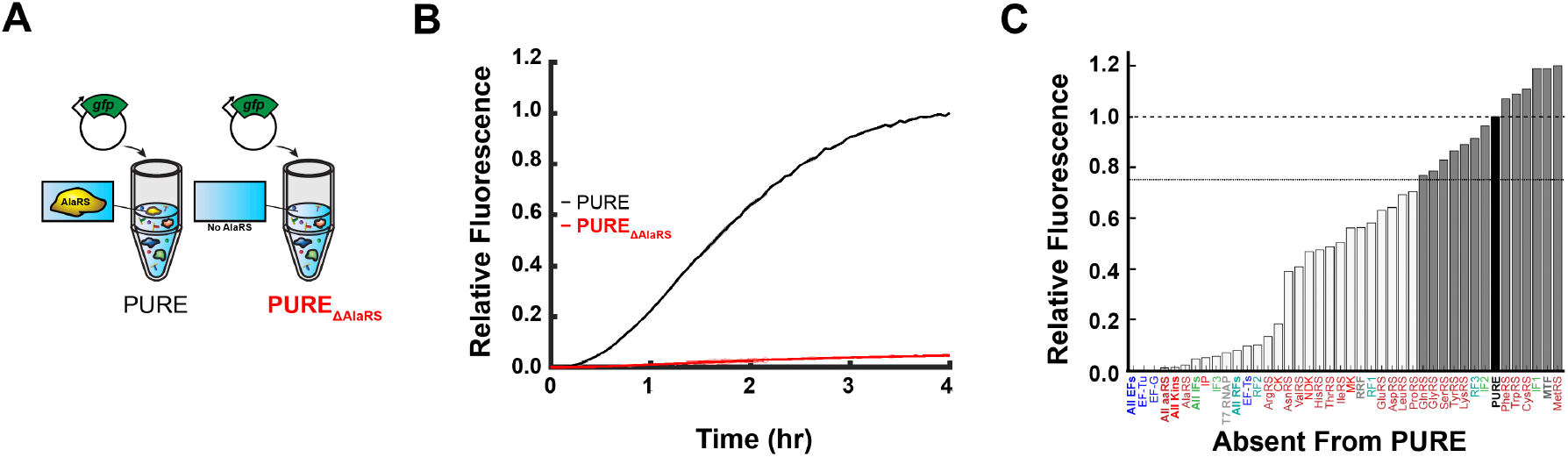
Molecular ensembles lacking individual components can be evaluated for reduced activity. **(a)** PURE lacking the alanine tRNA synthetase, PURE_△ (AlaRS)_, can be made and compared to complete PURE. **(b)** Expression of a green fluorescent protein (GFP) reveals that the absence of AlaRS (red) almost entirely eliminates PURE gene expression activity compared to intact PURE (black). **(c)** Single-component depletions (SCDs) of PURE result in a range of expression levels. All expression values taken from four-hour time points of triplicate PURE reactions and normalized to complete PURE (black bar, top dashed line). SCDs are rank ordered from lowest-to-highest expression level (left to right). Most SCDs produced enough expression reduction to support reconstitution assays (light grey bars, 75% or lower expression threshold, lower dashed line). Some SCDs did not result in significant reduction in gene expression (dark grey bars). Time course curves and standard deviations for individual and groups of enzymes can be found in Figs. S6 and S7.

We used an expression-loss threshold of 75 percent full PURE activity to select which SCDs to advance for use in testing if PURE-made components can reconstitute PURE. We selected this threshold empirically to provide sufficient dynamic range in our subsequent reconstitution experiments; we expect that optimized reporters and improved measurement methods will be helpful evaluating the thirteen components we did not consider further here (nine aaRSs, IF1, IF2, MTF, and RF3). The remaining 23 SCDs, each exhibiting a sufficient decrease in reporter gene expression, include all translation elongation factors, all PURE-specific kinases, T7 RNA polymerase (RNAP), and eleven aaRSs.

We next used PURE to produce PURE-made versions for each of the remaining 23 components. Specifically, we made expression vectors for each component, to be used with PURE, adding a twin-strep tag to the N-terminus of each component. We then expressed each component in PURE, as opposed to *E. coli*, performed strep-based affinity purification to recover the PURE-made component (PMC), and used SDS-PAGE to verify the mass and purity of each preparation (e.g., AlaRS, Fig. 3AB). We successfully expressed, purified, and verified the mass for 21 of the 23 PMCs. Ribosome Release Factor (RRF) is a small protein (2.98 kDa) and was difficult to confirm via SDS-PAGE gels, although we used the purified eluent for subsequent reconstitution experiments. We failed to obtain verified-mass T7 RNAP by our method.

**Figure 3.**
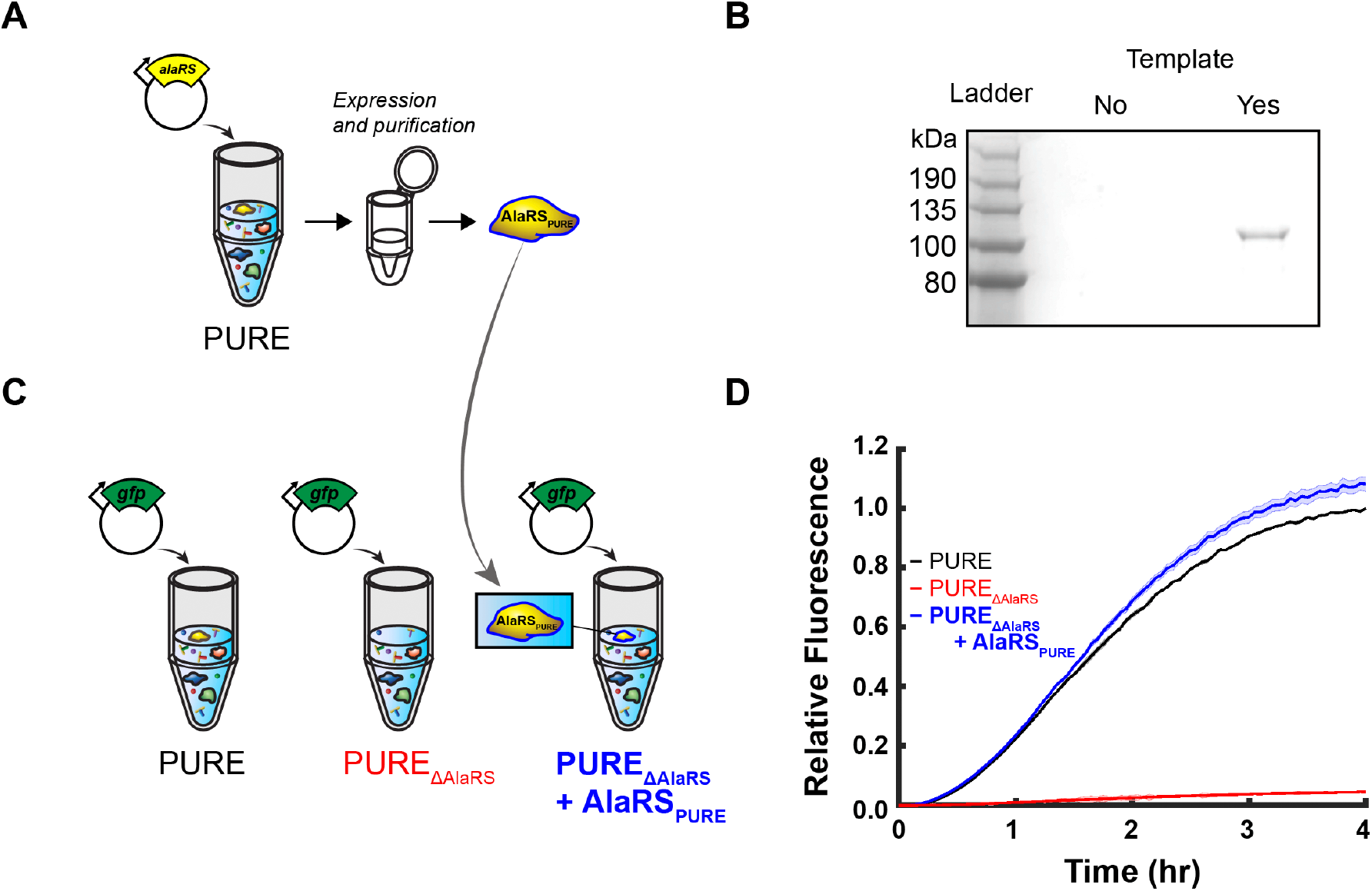
A molecular ensemble lacking an individual component can be complemented with an ensemble-made component. **(a)** As one representative example, an expression vector for N-terminal dual strep-tagged AlaRS is added to PURE. **(b)** PURE-expressed AlaRS is purified and validated for mass and homogeneity via SDS-PAGE and staining. **(c)** PURE-made AlaRS is added to PURE lacking AlaRS, PURE_△ (AlaRS)_, to evaluate whether PURE-made AlaRS is functional. **(d)** Expression time series for a GFP reporter in PURE_△ (AlaRS)_ (red), PURE_△ (AlaRS)_complemented with *E. coli*-made AlaRS (black), and PURE_△ (AlaRS)_ complemented with PURE-made AlaRS (blue).

We then performed complementation experiments in which we supplemented each SCD with its cognate PMC and observed if and to what extent PURE activity was recovered. As independent benchmarks, we also supplemented each SCD with the missing component as made by expression and purification from *E. coli* cells (i.e., *E. coli*-made components, or EcMCs). We added the same component final concentrations for both the PMC- and EcMC-complementation reactions, allowing us to more directly discriminate between any differences in expression arising via differences in functional activity of components due to source (i.e., PURE or intact cells), versus differences in expression arising merely due to differences in the concentration of each component in the supplemented SCD reactions relative to commercial PURE. As one example, we added equal amounts of PMC- and EcMC-produced AlaRS to two independent PURE_△(AlaRS)_samples and observed GFP expression (Fig. 3C), finding that both mixtures produced nearly-equal and high levels of expression, consistent with commercial PURE (Fig.3D), suggesting that functional AlaRS can be remade by PURE alone. We performed similar experiments for the 21 remaining SCDs and observed differing levels of expression recovery for each (Figs. S5-6, blue curves).

To represent if and to what extent a system can remake itself at a single-component level we developed standard quantitative metrics that can be used for any component whose function can be transduced to expression of a reporter gene. Specifically, we used the levels of GFP expression from each SCD and PMC-complemented SCD relative to the GFP expression level obtained via intact PURE to define depletion and recovery scores, respectively (Fig. 4A). We used a split-box template and color map to visualize both values (Fig. 4B). We used this method of representation to abstract and quickly summarize the capacity of our assay to detect depleted components, as well as the capacity of the system to remake its individual components. As one example, depletion of Initiation Factor 2 (IF2) did not result in sufficient reduction of expression to warrant further study here (Fig. 4C). As a second example, depletion of HisRS reduced GFP expression to ∼0.47 of intact PURE, and subsequent complementation recovered ∼0.86 of intact levels (Fig. 4D). We observed that the expression dynamics across all experiments followed a pattern, with maximum observed expression by four hours; thus, all values represented here were calculated from four-hour time points.

**Figure 4.**
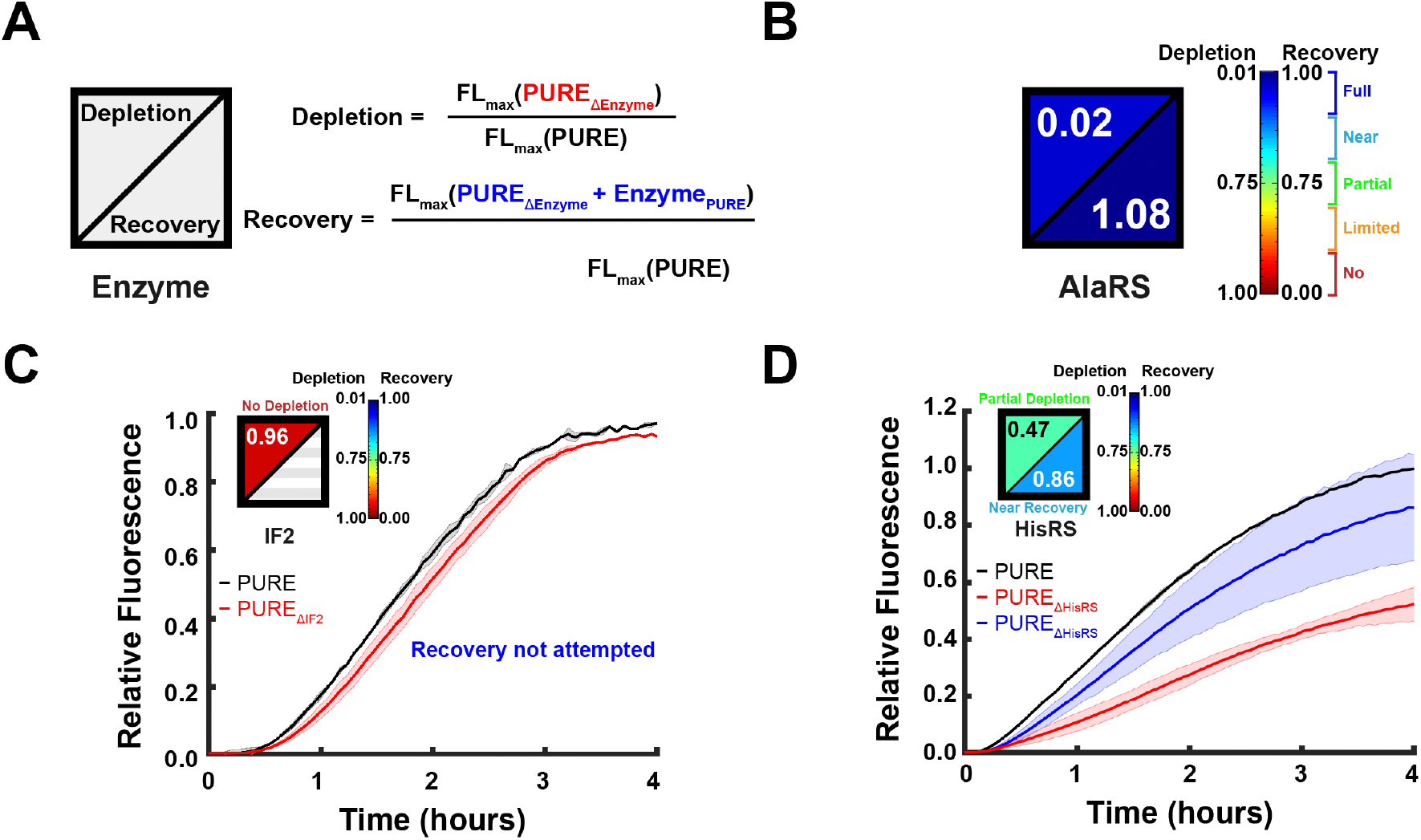
A standard and quantitative metric can be used to evaluate the extent to which an ensemble loses and restores activity as components are removed and regenerated. **(a)** Depleted system activity is defined by the activity of the depleted system relative to the intact system. The regenerative capacity of the system is defined by the extent to which the system recovers full system activity when a depleted system is complemented with the missing system-made components. **(b)** Color maps selected to represent effects of depletion and complementation on a single scale; depletion of AlaRS results in near full loss of activity, and complementation of PURE-made AlaRS recovers full system activity. **(c)** Representative example for a component, Initiation Factor 1 (IF1) whose individual depletion does not result in significant reduction of system activity. **(d)** Representative example for a component, Histidine tRNA synthetase (HisRS), whose individual depletion results in a partial reduction of system activity, and complementation recovers near-full system activity.

Inspired by the utility of the Periodic Table in organizing and representing humanity’s collective understanding of the fundamental chemical elements comprising matter, we explored if and how to best organize our representations of the capacities of fundamental life-essential functions. We grouped the split-box for each component into higher-order life-essential clusters (e.g.,transcription, aminoacylation, recycling of tRNA, ribosomes, energy carriers, and translation factors). Because we choose to study the PURE system here, as have others, as a foundation from which to support ongoing bottom-up synthetic cell building efforts, we named our integrated visual representation the “Pureiodic Table” (Fig. 5). Going forward, as still-more life-essential and life-contributing functions are added and evaluated, we imagine that “pure” will refer only to the fact that purified components are being tested within the context of a well-defined ensemble of molecules.

**Figure 5.**
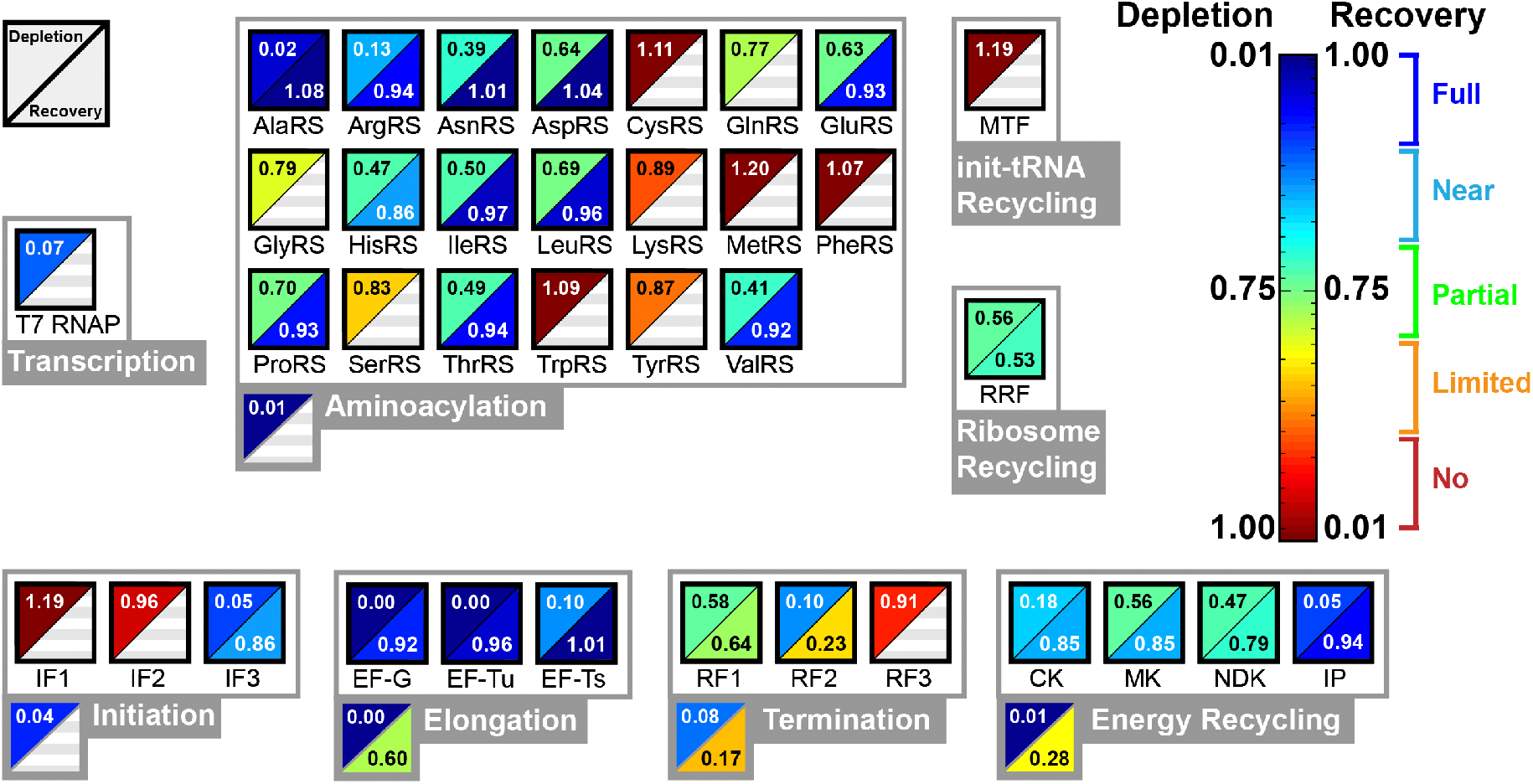
A Pureiodic Table representing the capacity for functional regeneration among the 36 enzymes in PURE. Quantitative depletion and recovery scores for all individual components tested. Some components did not result in sufficient loss of system activity when depleted to warrant complementation testing (lower right triangles with grey stripes). Components are clustered on basis of higher-order life-essential functions (as-labeled grey box outlines). Cluster-wide depletion and recovery assays (i.e., removing and restoring all cluster- specific components at once) reveal intra-cluster functional complementation hidden by single-component tests.

Practically, from the initial table, we can quickly discern that the blue/blue boxes represent components that have the greatest certainty of functional auto-regeneration, including AlaRS, AergRS, EF-G, EF-Ts, ET-Tu, IF3, CK (creatine kinase), and IP (inorganic pyrophosphatase). Perhaps more importantly, we can also quickly identify components that still need validation (red/empty boxes) as well as components that cannot now be sufficiently regenerated. For example, PURE-made RRF, RF1, and RF2 did not recover PURE-based gene expression in PMD-complemented SCDs. Finally, in addition to presenting the properties at a single-component level, we also created cluster-wide boxes for subsets of functions that we could perform cluster-wide depletion and complementation experiments. For example, each individual translation elongation factor could be removed and fully regenerated as measured by our individual assays but, when all three components were depleted and complemented together, only ∼0.60 full PURE activity was recovered, suggesting that redundant activities within this trio mask partial loss of function in the SCD-based assays.

Well aware that realizing fully-reproducing synthetic cells is expected to involve integration and testing of hundreds of components, and that others are pursuing and reporting on the regenerative capacities of molecular systems including PURE, we sought to explore integration of reported results across groups in the form of an expanded Pureiodic Table. To do so we incorporated data, where available from the literature, for the components tested here as well as for tRNA and ribosome regeneration (Fig. 6). While it is immediately apparent that much work remains, a simple and standard visual representation that can be simply shared and quickly updated should facilitate collaboration and support the scaling of team-based efforts seeking to accelerate synthetic cell building.

**Figure 6.**
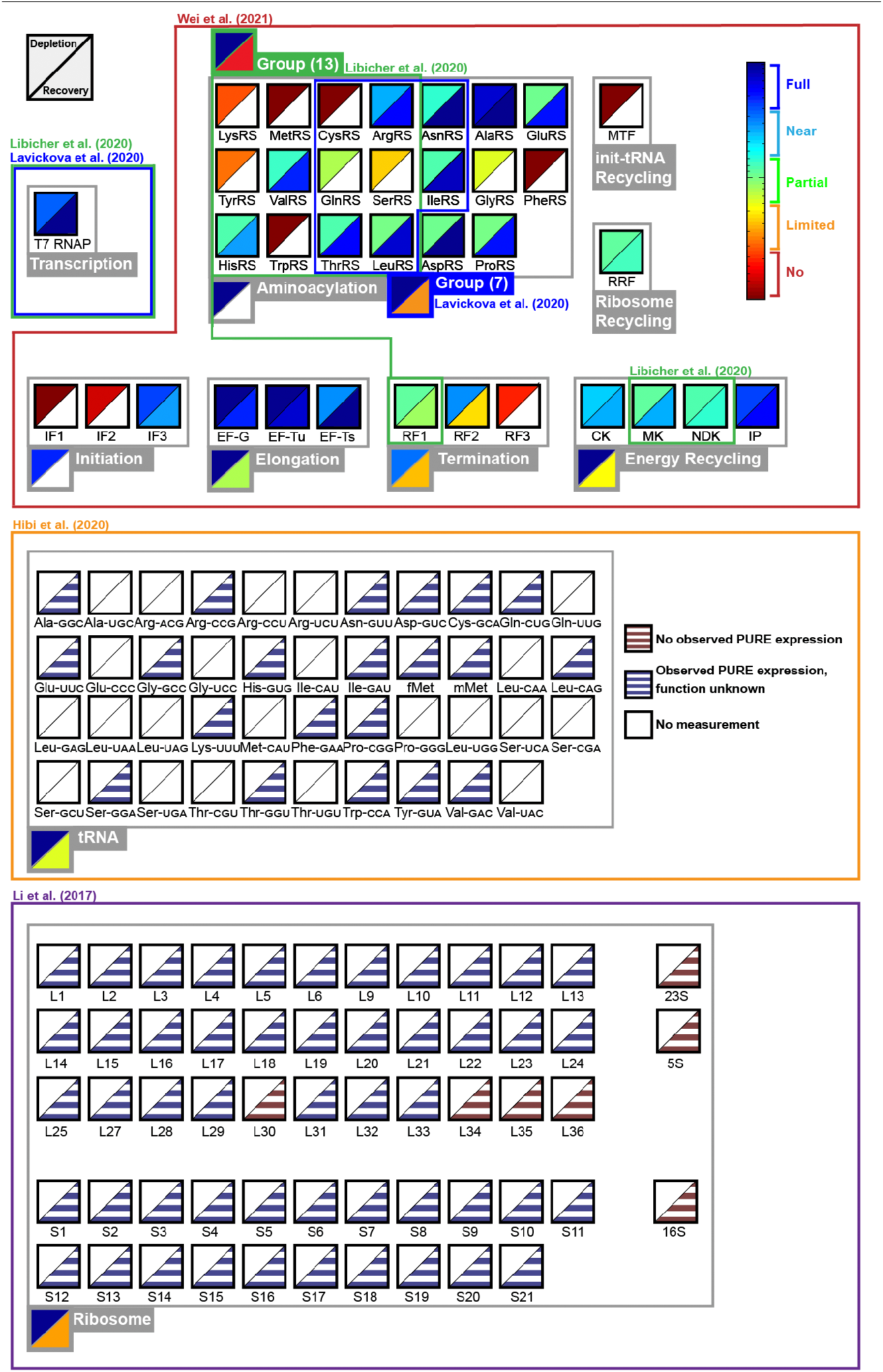
An expanded Pureiodic Table represents community-wide results and enables coordination of effort towards synthetic cell building. The table as in Fig. 5 but with overlapping literature data incorporated as possible. Colored-block outlines indicate specific sources of data and regions of overlap. Additional life-essential components (e.g., tRNA, ribosomes) are incorporated as possible. Empty blocks and triangles (white) highlight individual components for which data is not yet available. Data referenced from [27-30].

### Discussion

We tested if PURE can express functional versions all of the 36 single-enzyme components comprising PURE. We constructed single-enzyme dropout PURE reactions and measured the resulting decreases in reporter gene expression. 23 of the 36 enzymes tested, when absent from PURE, resulted in gene expression levels below 75 percent of full PURE. We then used PURE to remake these 23 enzymes and successfully purified 21 of the 23 attempted. We added each enzyme back to its cognate dropout PURE and measured any recovery in gene expression levels. 19 of the 22 enzymes tested recovered some or all of full PURE activity. We developed a standard quantitative visual framework for representing our results, the Pureiodic Table, as a tool for enabling community formation and fellowship among and within synthetic cell building efforts.

The PURE system is represented to be a minimal transcription and translation system, in the sense that expression should decrease – at least 50% reduction across all enzymes and over 90% reduction for 20 enzymes – if any single component is removed from the system [16]. Instead we found that 13 of the 36 individual dropouts tested here did not result in significant reductions in gene expression. One difference is our use of GFP as a reporter versus DHFR in the original study and any resulting differences in demand for individual amino acids (Table S1).

Development and testing of reporters optimized for probing the reproductive capacity of PURE, and of genetic devices that transduce other biochemical activities into gene expression signals, can improve and expand the scope of future testing efforts.

We found that three enzymes – RRF, RF1, and RF2 – did not result in recovered PURE activity when tested. As noted, we could not verify production of RRF via SDS PAGE due to its small size even though we were able to detect in the purified sample a 0.81 ratio of 260 nm/280 nm via spectroscopy. Thus, we cannot now discriminate between the possible absence of RRF in our complementation test versus the possibility that PURE-made RRF is non-functional, either due to a missing modification when made by PURE or due to the presence of a N-terminal strep tag.

We suspect that the failure of PURE-made RF1 and RF2 in recovering gene expression are due to the likely absence of methylation. Mora et al. previously reported that RF1 and RF2 activities are greatly reduced if a universally conserved tripeptide, GGQ, is not methylated [33]. The required methylase, while present in the *E. coli* used to manufacture PURE, is not included in PURE thereby hindering the qualitative ability of PURE to support functional self-reproduction. Complementing PURE with the needed methylation function is an obvious next step that should also include additional tests that ensure the so-added methylase can be remade via the expanded PURE.

We note that while many PURE functions could be individually removed and restored via PURE-made copies, several groupings of functions performed differently when tested together. For example, IF1 or IF2 could be individually removed with no impact on gene expression but could not be removed in combination with IF3. As a second example, all three elongation factors could be individually removed and replaced with near total loss and full recovery of gene expression. However, when all three were simultaneously replaced with PURE-made versions gene expression only recovered to ∼60 percent of *E. coli*-made PURE levels. As a third example, the four energy recycling factors only recovered ∼40 percent of *E. coli*-made PURE levels when replaced all together. The difference between individual and ensemble recovery levels may be due to compounding impacts of partially-functional and complementary components. For example, no individual kinase showed a recovery of less than 79 percent yet the recovery via the PURE-made ‘energy recycling’ ensemble was only 28 percent. If each kinase was entirely functionally independent of the others we would expect 54 percent recovery across the ensemble (0.85*0.85*0.79*0.94), suggesting that among these four components there may be some functional complementation occurring in the single-component assays.

A simpler-to-setup assay that avoids the need to purify PURE-expressed enzymes involves providing the template encoding the component(s) of interest directly into the starting PURE mixture. So long as there is sufficient PURE activity in the starting mixture to kickstart the process, the PURE mixture itself remakes the partially-depleted component. We explored this approach using AlaRS as a test case, first observing GFP expression dynamics in PURE containing various initial concentrations of AlaRS, with and without DNA expressing AlaRS (Fig. S1). However, the challenge with this type of experiment lies in carefully discerning PURE activity arising due to the PURE-expressed enzymes versus enzymes present in the starting PURE, sourced from *E. coli*. To address this challenge Libicher et al. [30] proposed a serial transfer technique, whereby the initial concentration of the target components is increasingly minimized. Discrete depletion and complementation assays have the advantage of eliminating any crosstalk from the original non-PURE expressed enzymes and offer a simpler readout. However, fractional depletion and dilution approaches may serve to improve dynamic range for some components and enable for more widespread testing.

We hope that the expanded Pureiodic Table (Fig. 6) will serve as a framework for tracking humanity’s collective capacity to construct life from scratch. Many additional life-essential functions remain to be added and tested (e.g., replication, metabolism, membrane formation, cytokinesis, etc.). Each life-essential function so-represented serves as a visual icon of what is required to support autonomous reproduction. We also expect that factors improving ensemble performance, including dynamic control and expression fine-tuning, will be required and should be represented. For example, the addition of chaperones and Elongation Factor P (EF-P) to PURE has been shown to increase protein yield and quality [17]. There may also be chemical or physical aspects essential to cell building that are not directly genetically encoded. As the Pureiodic Table is expanded we look forward to collectively confronting a seemingly complete table and pondering, if needed, what life-essential unknowns remain to be discovered.

### Materials and Methods

#### Plasmids and strains

All plasmids used in this work will be made freely available as the “Pureiodic Table Construction Kit” via the bionet node maintained by the Stanford Freegene project (stanford.freegenes.org) and, independently, via Addgene (addgene.org). We used a pET-28a vector and *E. coli* BL21(DE3) for all 36 single-enzyme expression preparations following established protocols [34]. We used a pPSG-IBA105 vector (IBA Lifesciences), including a T7 RNAP promoter and twin strep tag [35], to clone and express enzymes in PURE. We modified the iGEM pSB3K5 (BBa_I50032) by replacing the promoter with a T7 RNAP promoter and adding sfGFP to its open reading frame.

#### PURE cell-free expression

We purchased all commercial PURE-related materials from New England Biolabs (neb.com). Each reaction was constructed from 5 ul of Solution A (NEB), 3.5 ul of Solution B (NEB or made from scratch as reported here), 1.25 ul of DNA template (5 nM final concentration), 1.25 ul of RNAse inhibitor, Murine (NEB), and 1.25 ul water or as reported. For complementation reactions we matched the final enzyme concentration as reported in the academic literature [36]. The reaction was pipette-mixed and a 10 µl aliquot of the total mixture is transferred to a 384- well round clear-bottom plate (Corning). Reactions were mixed and initialized at 4C in a cold room, transferred to plates, and then sealed with clear micro-plate tape (Corning) before being placed in a plate reader (SpectraMax i3) at 37C. We obtained measurements every three minutes over four hours. We used 485 nm excitation (10 nm bandwidth) and 520 nm emission (10 nm bandwidth) wavelengths for measuring GFP levels.

#### Protein purification from E. coli

For His-tag purification of *E. coli* proteins we back-diluted 1:1000 an overnight culture into 2 L of LB. Following 3 hours of growth at 30C we added IPTG to 0.1 mM final concentration. After another 9-15 hours of growth, we pelleted cell cultures via centrifugation at 5000 x g at 4C for 20 minutes (JA-10 rotor). We responded cell pellets in 25 ml of equilibration buffer in 50 ml falcon tubes and sonicated four times using a microtip (duty cycle 50%, 45 seconds treatment with 2 minutes on ice in between treatments). We centrifuged the resulting solution at 15000 x g at 4C for 60 minutes (JA-10 Fixed-Angle Rotor, Beckman Coulter). We placed Ni-NTA slurry (Thermofisher) in columns with 20 ml of equilibration buffer running through followed by the sample-containing supernatant followed by 20 ml of wash buffer and 5 ml of elution buffer. We then ran the eluted mixture through a FPLC (AKTA PURE) with a salt gradient consisting of mixtures of buffer A (50 mM HEPES) and buffer B (50 mM HEPES plus 1M sodium chloride). We combined protein fractions, exchanged the buffer for storage (Amicon filter), and stored the so-purified proteins at -80C following flash freezing.

#### Protein purification from PURE

We added T7-expression vectors encoding strep-tagged proteins (5 nM concentration) in 200 ul of total PURE (NEB) working volume. We pre-washed 500 ul of strep-tactin beads (IBA Lifesciences) three times using 1 mL of wash buffer (Buffer W, IBA Lifescences) before adding the entire PURE reaction volume and incubating at 4C for 3 hours. We immobilized the magnetic beads using a magnet and washed (5x) via pipetting 1 ml of wash buffer. We added elution buffer (IBA Lifesciences) and incubated at 4C for 10 minutes. We then immobilized the bead while taking the supernatant, exchanged buffers for storage (Amicon filter), verified quality via gel separation and mass via a spectrophotometer (Nanodrop). We stored the proteins at -80C following flash freezing.

#### Buffer recipes

Our equilibration buffer consisted of 50 mM sodium phosphate, 300 mM sodium chloride, and 10 mM imidazole. Our wash buffer consisted of 300 mM sodium choloride, and 25 mM amidazole. Our elution buffer consisted of 300 mM sodium chloride and 250 mM imidazole. Our regeneration buffer consisted of 20 mM MES sodium and 100 mM sodium chloride. Our protein storage buffer consisted of 50 mM HEPES, 100 mM potassium chloride, 10 mM magnesium chloride, 7 mM 2-mercaptoethanol, and 30% glycerol.

#### Protein gels

We ran our SDS-PAGE separations with 10% Bis-Tris in MOPS buffer, or MES buffer for smaller proteins (∼10 kDA), typically at 200 volts for 30 minutes followed by staining with Instant Blue (Radeon) for 15 minutes.

#### Quantitative estimate of peptide-bond formation capacity of batch PURE

Adding up the total concentrations of proteins in PURE [36], excluding ribosomes, yields ∼5,400 µg/mL or ∼3*10^7^ peptide bonds/fL; Including ribosomal proteins increases the total PURE protein concentration to ∼17,400 µg/mL or ∼10^9^ peptide bonds/fL. Meanwhile, the expression capacity of PURE is ∼200 µg/mL or ∼10^6^ peptide bonds/fL [NEB; confirmed experimentally]. Thus, the capacity of PURE to synthesize proteins would need to be increased ∼27-fold, or ∼87-fold including ribosomal proteins, to satisfy the quantitative reproduction requirement (Fig. 1C).

## Acknowledgements

We thank Conary Meyer for discussions and advice in formulating and framing the work presented here. We thank CM and Alec Lourenço for helping with the cloning of pET constructs. We thank Corinna Tuckey and New England BioLabs for constructing and providing customized PURE kits. We thank Kai Yuet, Ray Zhuang, James Payne, and Tim Valentic for guidance with protein purification and access to FPLC. We thank the Endy Lab for discussions and general support. Funding was provided, in part, by a U.S. National Science Foundation Graduate Fellowship and a Stanford Graduate Fellowship to EW.

---

**Supplementary Figure 1.**
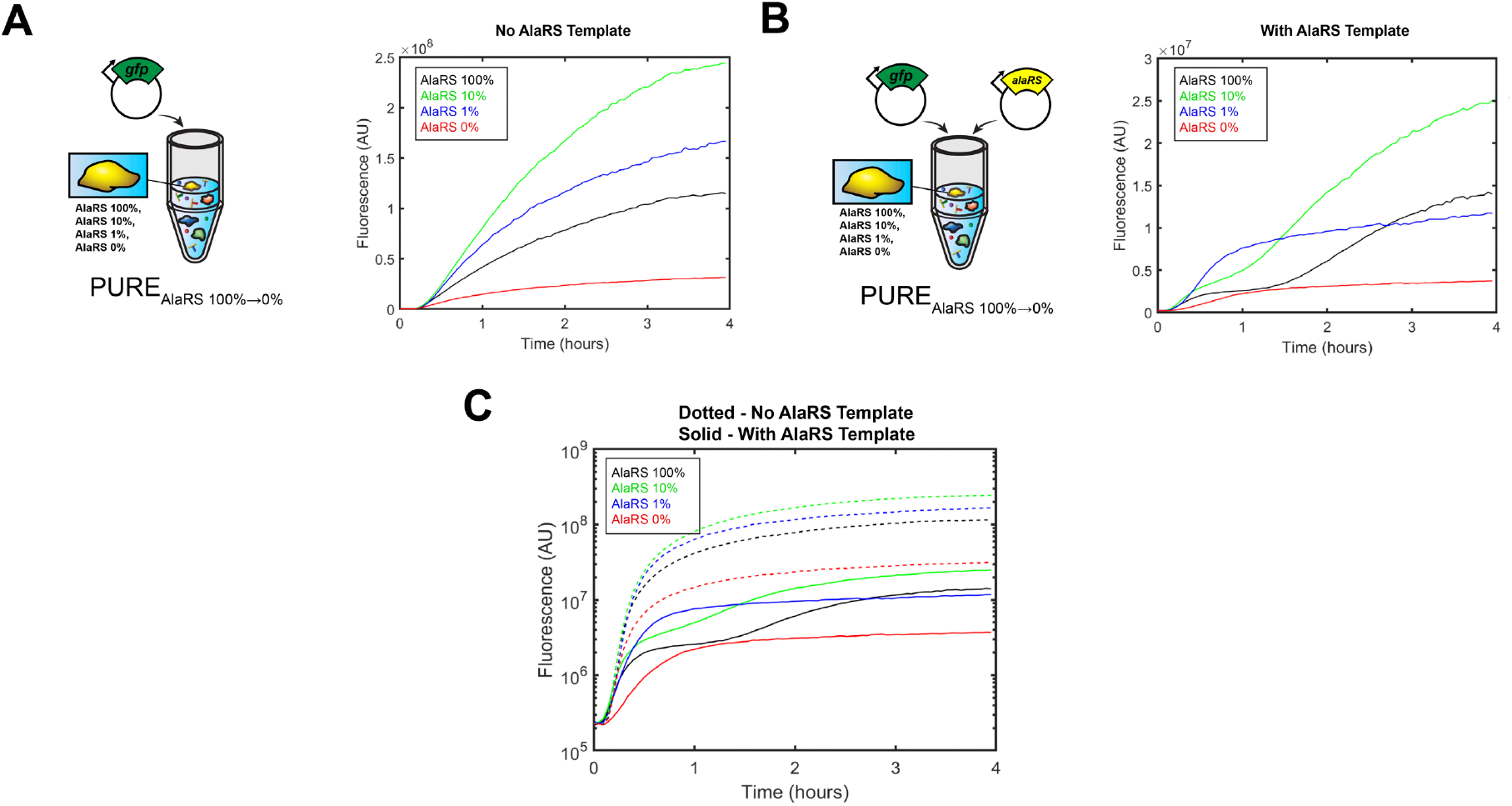
Varying the concentration of a PURE-essential component impacts report gene expression, and providing DNA template expressing the same component impacts expression dynamics and levels. **(a)** PURE lacking AlaRS was complemented with a range of initial concentrations of AlaRS (0%, 1%, 10%, 100% PURE) and expression of a GFP reporter (2.5 nM DNA concentration) was measured. **(b)** When a DNA template expressing AlaRS is added to the initial system (2.5 nM DNA concentration), GFP expression dynamics change and decrease. **(c)** data from **(a)** and **(b)** plotted on a log10 y-axis.

**Supplementary Figure 2.**
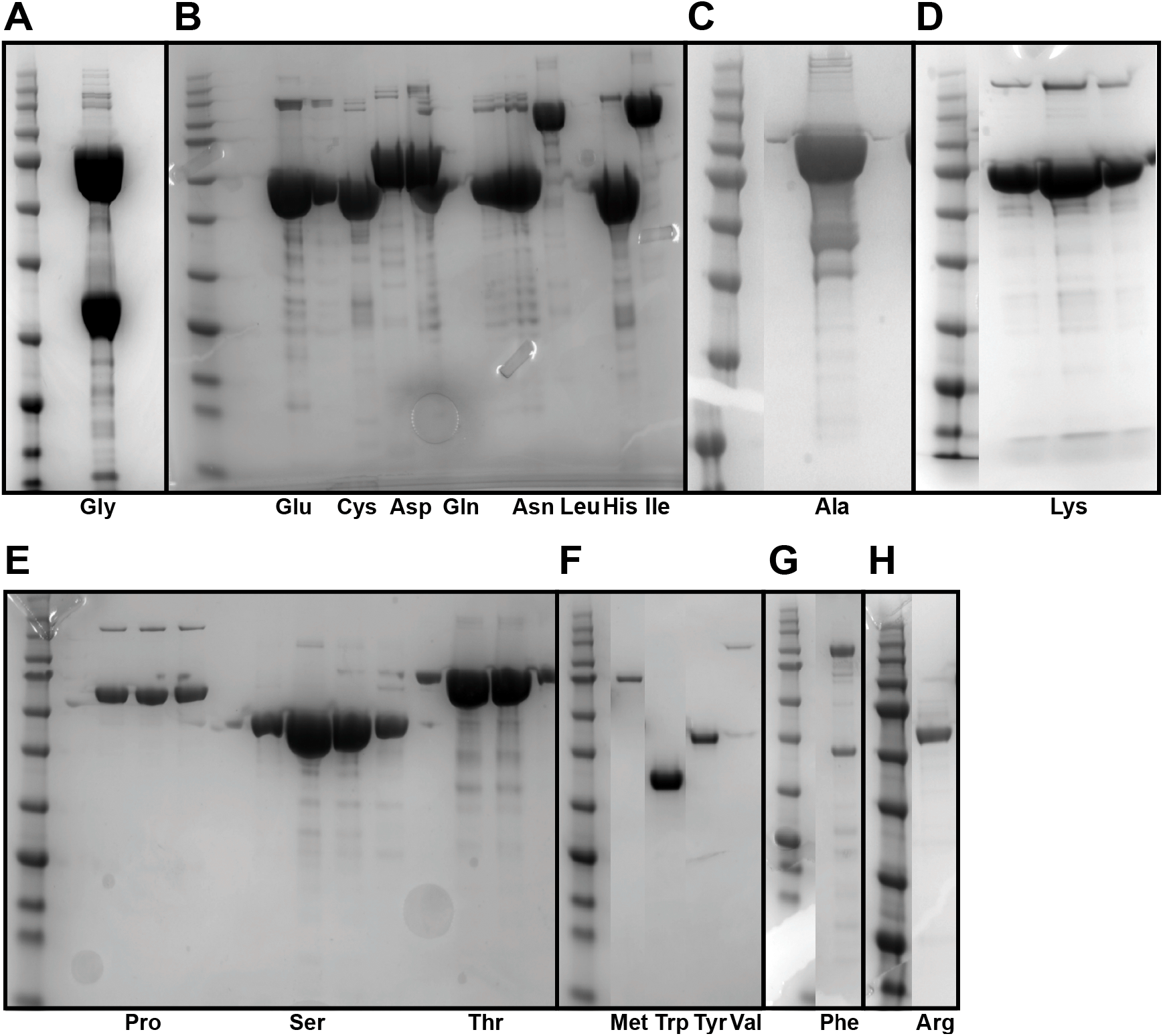
Production and purification of all 20 *E. coli* tRNA synthetases from *E. coli*. **(A-H)** Individual tRNA synthetases as indicated below each experimental lane. A broad range pre-stained protein standard (NEB #P7712) was used as a ladder. Each individual protein was His-purified using a Ni-NTA resin column along with FPLC and buffer exchange to increase purity. Expression, purification, gel, and buffer details as described (Materials and Methods).

**Supplementary Figure 3.**
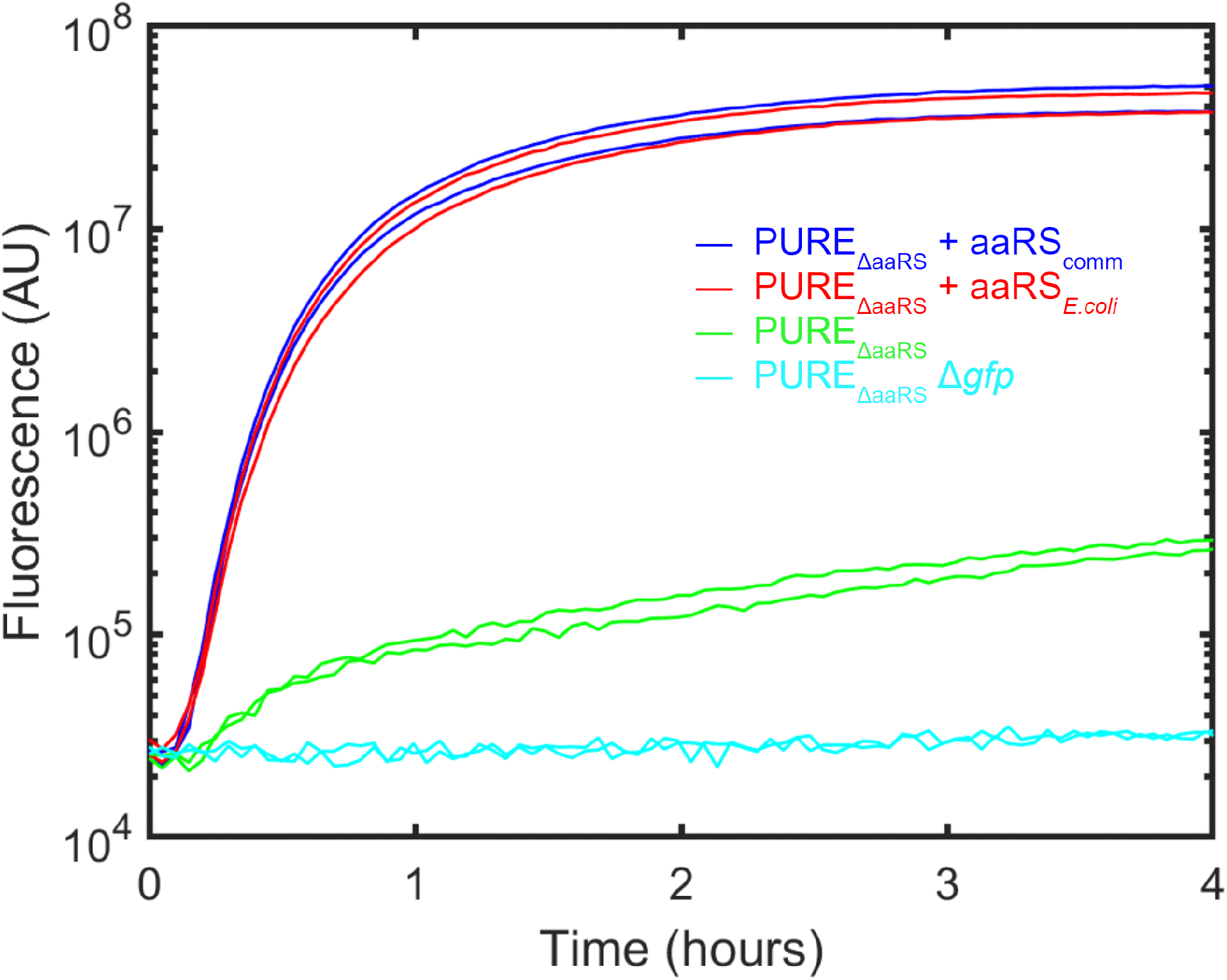
Commercially-sourced and locally-made tRNA synthetases both restore expression activity. Expression of a GFP reporter in PURE lacking a GFP expression template (light blue), containing a GFP expression template but lacking all tRNA synthetases (bright green), and containing a GFP expression template plus tRNA synthetases either sourced commercially (blue) or made locally via purification from *E. coli* (red).

**Supplementary Figure 4.**
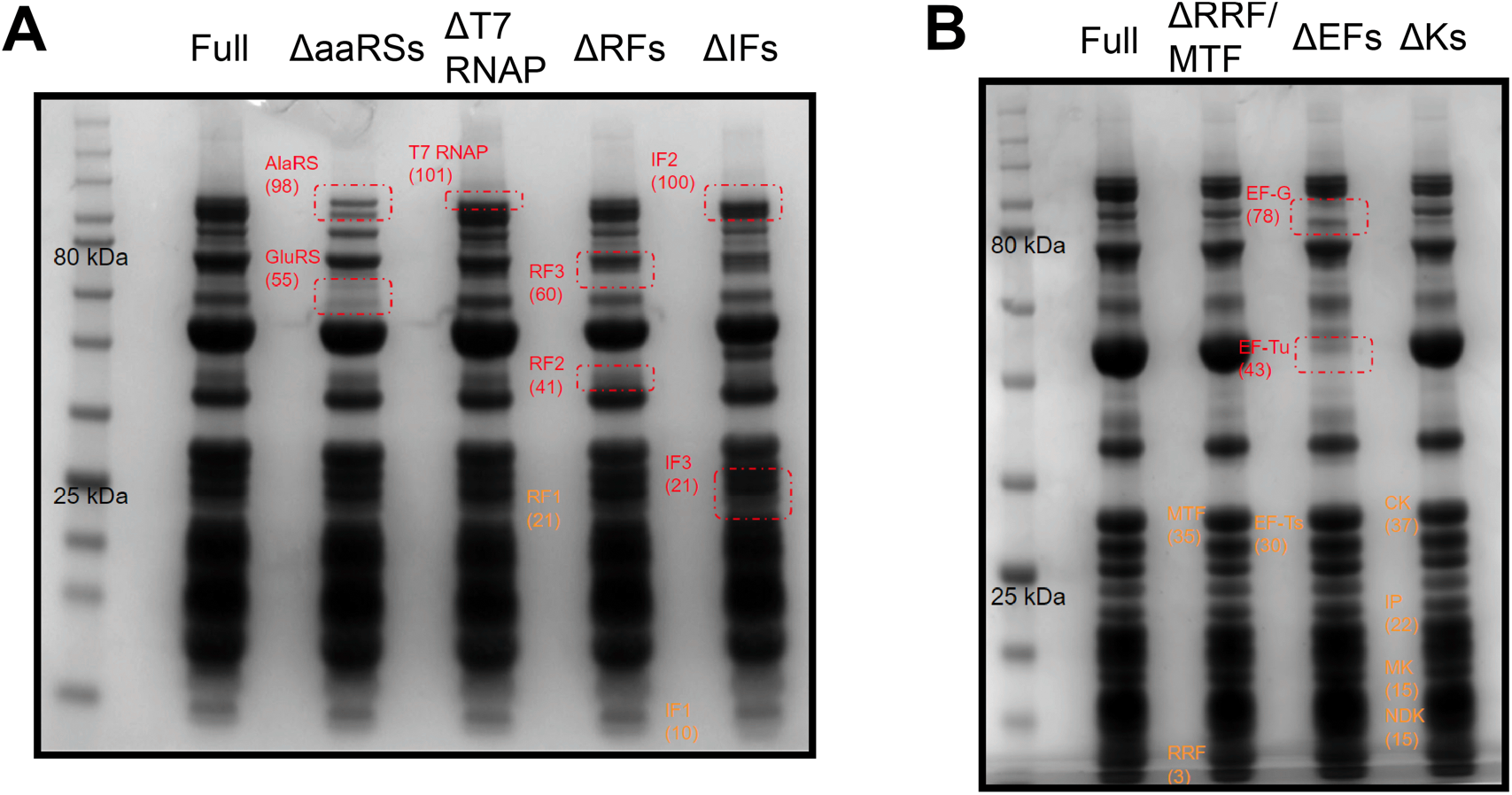
Verification of dropout PURE mixtures via 1-D protein gel electrophoresis. SDS-PAGE results for each custom dropout Solution B (NEB). **(a)** from left to right, normal Solution B (full), missing all tRNA synthetases (△aaRSs), missing T7 RNA polymerase (△T7 RNAP), missing all release factors (ΔRFs), and missing all initiation factors (△IFs). **(b)** from left to right, normal Solution B (full), missing ribosome and tRNA recycling factors (△RRF/MTF), missing elongation factors (ΔEFs), and missing energy recycling factors (△Ks). A broad range pre-stained protein standard (NEB #P7712) was used as a ladder for each gel. Expected location of specific components as noted (red, orange).

**Supplementary Figure 5.**
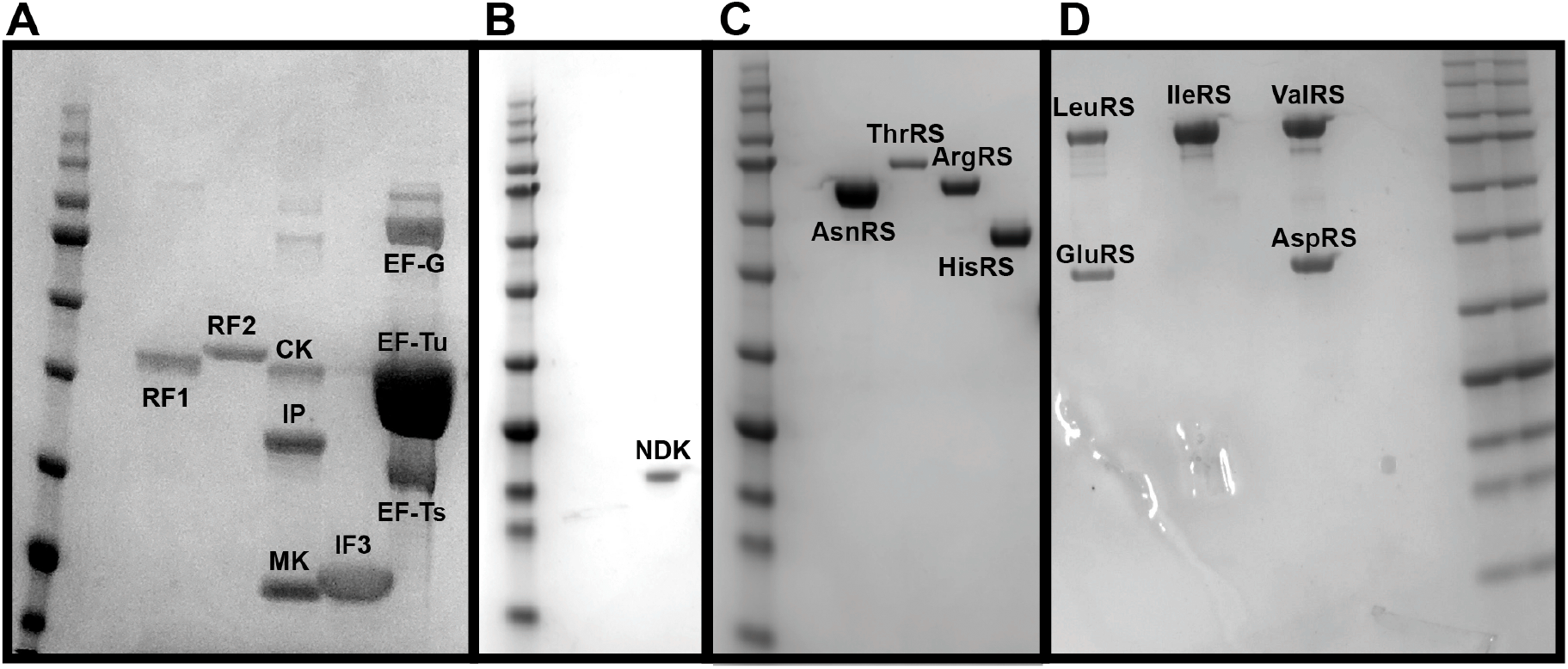
Production and purification of PURE components from PURE. **(a-d)** various PURE components made via expression and purification from PURE as describe (Materials and Methods) and noted directly on each gel. A broad range pre-stained protein standard (NEB #P7712) was used as a ladder for each gel.

**Supplementary Figure 6.**
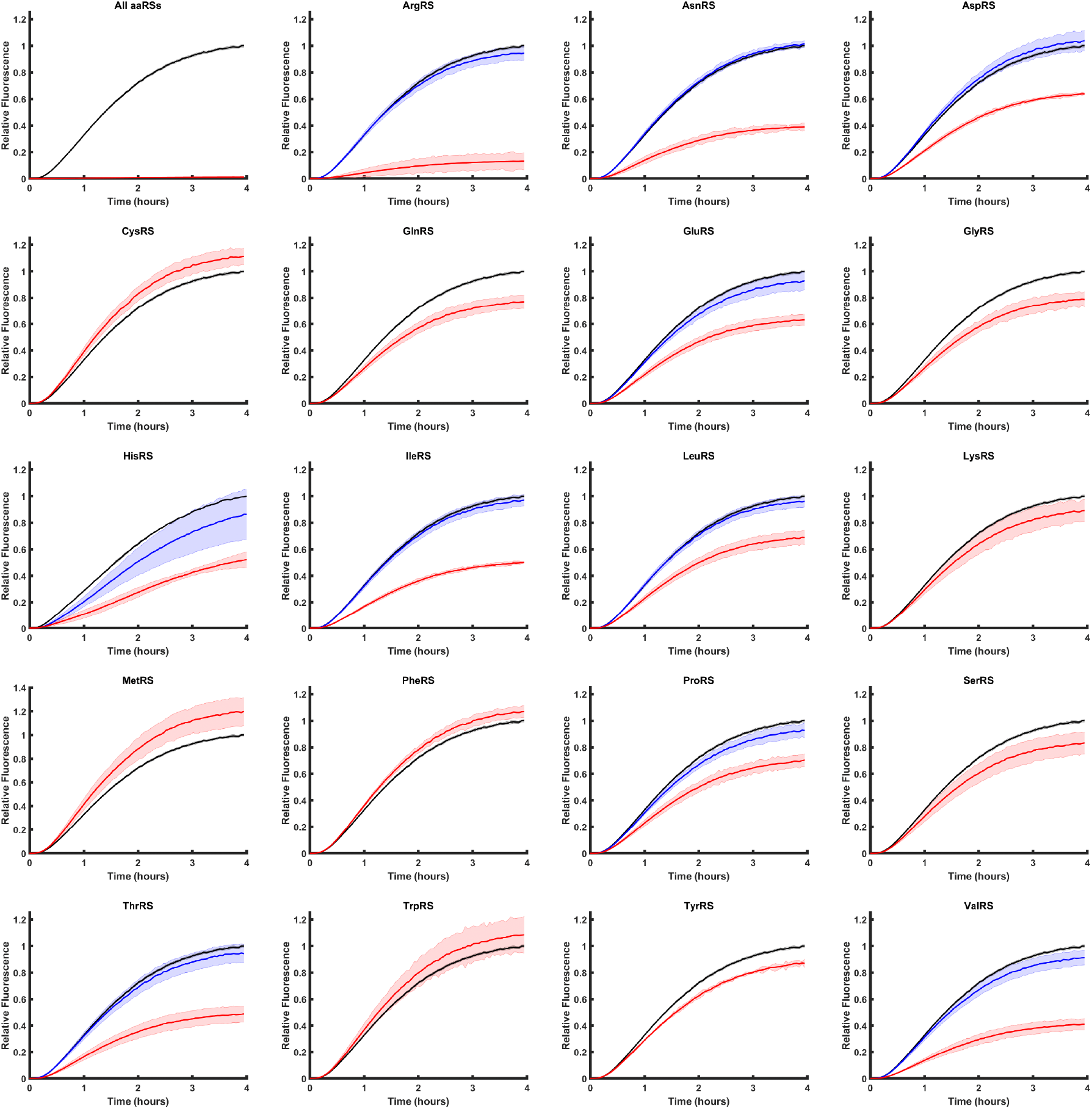
GFP expression profiles obtained as described (Materials and Methods) for all individual PURE depletion mixtures (as noted) and some combinations of PURE components (e.g., all aaRSs, all IFs, all EFs, all RFs, and all Ks). Shaded region indicated standard deviation in observed fluorescence for each specific experiment measured in triplicate. GFP expression trajectories were obtained for full PURE (black), depleted PURE (red), and, for depletions exhibiting below 75% full PURE activity, complemented PURE (blue). The values for each experiment were normalized to each experiment’s average full PURE value obtained at four hours.

**Supplementary Figure 7.**
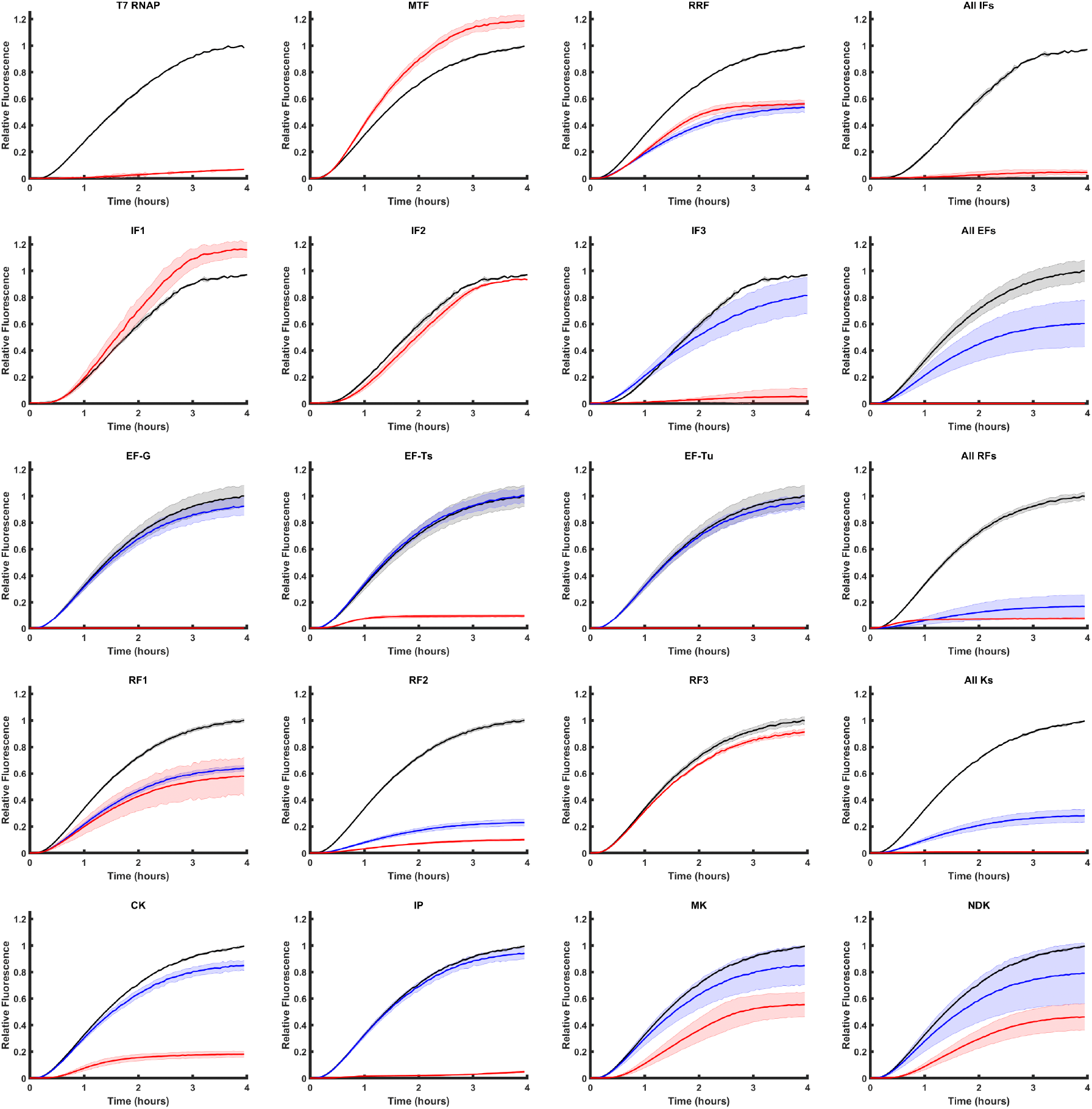
GFP expression profiles obtained as described (Materials and Methods) for all individual PURE depletion mixtures (as noted) and some combinations of PURE components (e.g., all aaRSs, all IFs, all EFs, all RFs, and all Ks). Shaded region indicated standard deviation in observed fluorescence for each specific experiment measured in triplicate. GFP expression trajectories were obtained for full PURE (black), depleted PURE (red), and, for depletions exhibiting below 75% full PURE activity, complemented PURE (blue). The values for each experiment were normalized to each experiment’s average full PURE value obtained at four hours.

**Supplementary Table 1.** A given reporter gene may have characteristics that affect suitability for measuring auto-regenerative systems. **(AA)** the twenty canonical amino acids with given three letter abbreviations.**(a)** the number of residues for each amino acid in the reporter gene used here (green fluorescent protein, GFP). **(c)** the concentration of each cognate amino acid’s tRNA synthetase in the PURE system. **(d)** the “demand” for a given amino acid normalized to the capacity of PURE to recharge spent tRNA. **(r)** the “regeneration number” as defined by the number of residues for each amino acid in its cognate tRNA synthetase. **(x)** the ratio “demand” to “regeneration number.”

| Amino Acid (AA) | a = AA # in GFP | c = aaRS concentration in PURE (mg/mL) | Load-balanced GFP expression demand, d = a/c | r = AA # in cognate aaRS | x = d/r = a/(rc) |
| --- | --- | --- | --- | --- | --- |
| Ala | 8 | 690 | 11.6 | 91 | 0.13 |
| Arg | 8 | 20 | 400 | 32 | 12.5 |
| Asn | 13 | 220 | 59.1 | 20 | 2.95 |
| Asp | 18 | 80 | 225 | 46 | 4.89 |
| Cys | 2 | 12 | 167 | 5 | 33.3 |
| Gln | 7 | 38 | 184 | 15 | 12.3 |
| Glu | 16 | 126 | 127 | 39 | 3.26 |
| Gly | 22 | 96 | 229 | 22 | 10.4 |
| His | 10 | 8 | 1250 | 8 | 156 |
| Ile | 11 | 400 | 27.5 | 46 | 0.60 |
| Leu | 20 | 40 | 500 | 65 | 7.69 |
| Lys | 20 | 64 | 313 | 24 | 13.0 |
| Met | 5 | 21 | 238 | 21 | 11.3 |
| Phe | 12 | 17 | 70.6 | 20 | 3.53 |
| Pro | 10 | 100 | 100 | 28 | 3.57 |
| Ser | 10 | 19 | 526 | 18 | 29.2 |
| Thr | 18 | 63 | 286 | 24 | 11.9 |
| Trp | 1 | 11 | 90.9 | 2 | 45.5 |
| Tyr | 9 | 6 | 1500 | 11 | 136 |
| Val | 18 | 18 | 1000 | 61 | 16.4 |

